# BetaH proteolysis unleashes an electrostatic-homing antibacterial polymorphic toxin

**DOI:** 10.1101/2025.07.24.666436

**Authors:** Daniel Mwangi, Maureen K. Thomason, Lauren M. Shull, Qing Tang, Joshua J. Woodward

**Affiliations:** Department of Microbiology, University of Washington, Seattle, WA 98109, USA; Department of Biology, University of Texas at Arlington, Arlington, TX 76019, USA

## Abstract

Contact-dependent or diffusible proteinaceous polymorphic toxin systems (PTSs) mediate widespread bacterial competition. While bioinformatic analyses have identified diverse PTSs across bacterial phyla, experimental validation in Gram-positive species remains limited. Here, we characterize a diffusible PTS encoded by the *Staphylococcus aureus* S8-Ntox35 locus. We demonstrate that this system mediates inter-genus antagonism against *Listeria monocytogenes* via a bactericidal, heat-labile protein, and that toxicity depends on extracellular cleavage of the BetaH domain by an S8 peptidase. This processed peptide resembles a cationic antimicrobial peptide (CAMP) and facilitates intoxication of target cells by the Ntox35 RNase domain. Target cell resistance is impacted by known CAMP defense pathways, including DltABCD and MprF, and experimental evolution identified the ABC transporter AnrAB as essential for intoxication. Unexpectedly, disruption of AnrAB abolished Ntox35 susceptibility, while simultaneously sensitizing cells to the proposed CAMP like activity of the processed BetaH domain. These findings reveal a novel mechanism of inter-genus antagonism among Firmicutes and establish a functional role for extracellular processing and ABC transporter-mediated susceptibility in PTS activity. Our work expands the known repertoire of diffusible toxins in Gram-positive bacteria and sets the foundation for broader ecological and mechanistic investigation of S8-PTS systems.

**Significance:** Polymorphic toxin systems (PTSs) are widely used by bacteria to inhibit competitors, but diffusible proteinaceous toxins have been largely characterized in Gram-negative species. Here, we identify and mechanistically characterize a diffusible PTS in *Staphylococcus aureus* that mediates inter-genus antagonism in the Firmicutes phylum. This system employs a secreted S8 peptidase to cleave a BetaH-toxin fusion, releasing a previously caged cationic amphipathic α-helix that facilitates membrane targeting and intoxication of susceptible cells. We further show that toxin activity requires the target cell ABC transporter AnrAB, revealing a novel route of entry and an evolutionary tradeoff between toxin susceptibility and antimicrobial resistance. Together, these findings uncover a new mode of bacterial competition and highlight a broadly distributed toxin system with ecological relevance across Firmicutes.

## Introduction

To reproduce, bacteria must colonize environments where they can tolerate abiotic stresses and access nutrients needed for growth. Access to and maintenance of such a niche is frequently challenged by the presence of genetically distinct organisms that share overlapping ecological and nutritional requirements. Competition between these bacterial populations in both natural ecosystems and within mammalian hosts is pervasive. This has driven the evolution of a wide array of antagonistic weaponry that restrict replication of rival strains and species, allowing the producing organism and its clonal kin to thrive (1). Polymorphic toxin systems (PTSs) are a particularly prominent molecular armament and can be broadly categorized into two functional classes (2, 3). Contact-dependent systems utilize specialized secretion pathways and surface-associated proteins, (e.g. T6/T7SS and CDI, respectively) that directly deliver toxic effectors into neighboring cells upon physical contact (2–6). Contact-independent systems deploy diffusible antibacterial agents, including both small molecule and proteinaceous antibacterials (e.g. Colicins and Umbrella particles) that act at a distance to suppress competitor viability (7, 8).

Both contact dependent and independent PTSs exhibit a modular, multidomain architecture. A proteinaceous toxin domain targeting essential cellular processes is associated with one or more additional domains responsible for targeting and/or delivery into competitor organisms. The modular fusion of diverse toxic effectors to conserved delivery domains enables functional diversification within these PTSs. While the toxin domains themselves are highly variable and span a broad distribution of antimicrobial activities—including peptidoglycan degradation, membrane disruption, and nucleic acid cleavage—the conserved delivery components tend to restrict host range, typically limiting antagonistic activity to members of the same species or genus (3, 9). To date, few diffusible antibacterial proteins have been identified, and such systems have not been previously reported in Firmicutes.

Recent bioinformatic mining strategies that integrate domain architecture and genomic context have markedly accelerated the identification of predicted PTSs, including diffusible proteinaceous toxins associated with novel targeting modules (3, 10). One such system, deemed the S8-PTS, is encoded by a three-gene locus comprised of (1) a secreted S8 family serine peptidase, (2) a secreted variable toxin fused to a conserved BetaH domain, and (3) a cytosol-retained immunity protein. Among the ∼3,800 bacterial genomes initially reported as harboring this locus, many belong to the phylum Firmicutes (10). Here, we provide experimental evidence that this locus encodes a bona fide diffusible proteinaceous toxin capable of mediating antagonism across genus boundaries—an activity that distinguishes it from previously described toxins and suggests a broader ecological role in competitive interactions within Firmicutes. We demonstrate that the S8 peptidase functions to release a caged amphipathic cationic peptide within the BetaH domain, which electrostatically targets the membrane of recipient bacteria. Finally, we identify a susceptibility locus in a target organism that is required for intoxication. Notably, disruption of the transporter encoded by this locus reveals an evolutionary tradeoff, whereby resistance to the proteinaceous toxin is accompanied by increased susceptibility to non-enzymatic toxicity of the PTS and increased sensitivity to a small-molecule antibiotic against which the transporter normally confers protection.

## Results

### The S. aureus S8-Ntox35 locus encodes an inter-genus toxin

As approximately 65% of organisms containing the S8-PTS locus were within the *Staphylococcus* genus (10), we selected *Staphylococcus aureus* as a model organism to investigate the function of this system. *S. aureus* strains encoding this locus possess a secreted S8 family peptidase, a BetaH domain fused to a toxin domain termed Ntox35, and a cytosol-retained immunity protein (Fig. 1A). Structural prediction of the Ntox35 domain, coupled with a similarity search against the Protein Data Bank (PDB), revealed predicted homology to a characterized RNase toxin from *Brucella abortus*, consistent with prior bioinformatic work indicating that Ntox35 adopts a canonical BECR fold composed of an N-terminal α-helix followed by a four-stranded β-sheet (Fig. 1B) (3, 11).

**Figure 1.**
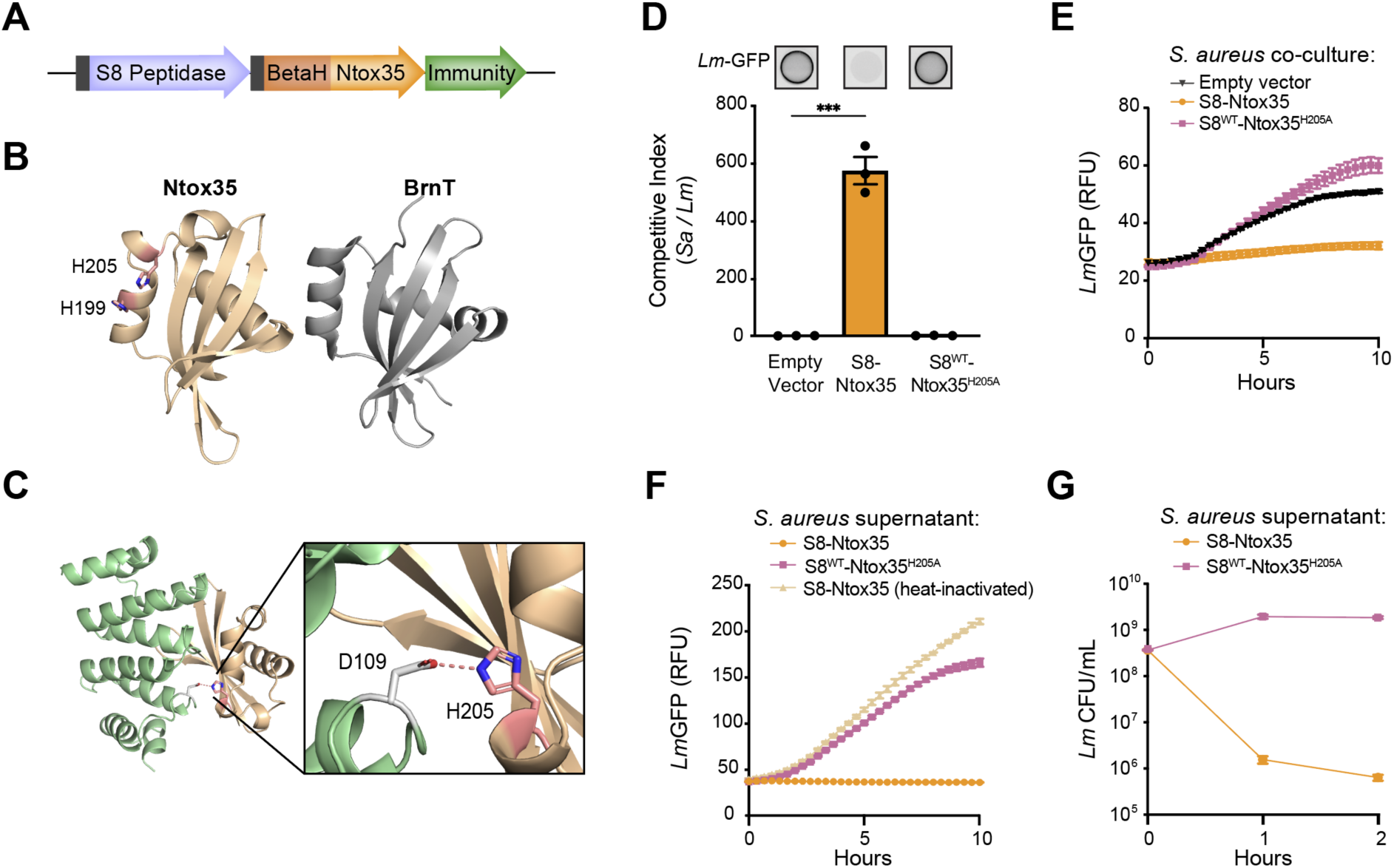
The S8-Ntox35 polymorphic toxin (PT) mediates diffusible interbacterial killing. (A) Three gene locus of the S8-PTS containing an S8 peptidase with a Sec signal sequence (grey box), a Sec signal containing BetaH domain fused to a variable toxin (Ntox35 for S. aureus), and an immunity protein. (B) Predicted structure of the Ntox35 toxin domain, generated using AlphaFold3, was queried using the Dali protein structure comparison server to identify a structurally related RNase toxin, BrnT (PDB: 3U97). (C) AlphaFold 3 structural modeling of the Ntox35 toxin domain (tan) and immunity protein (green) interaction. Enlarged inset shows the predicted interaction between Asp109 of the immunity protein and His205 of Ntox35. (D) Contact-dependent competition assay to evaluate the ability of *S. aureus* expressing either wild-type S8-Ntox35, catalytic Ntox35 mutant (S8^WT^-Ntox35^H205A^), or empty vector control to inhibit *L. monocytogenes* expressing GFP (*Lm*-GFP) on solid agar. Representative images show fluorescence of *Lm*-GFP in co-culture with the indicated *S. aureus* strain. Competitive indices were calculated as the ratio of *S. aureus* to *Lm*-GFP CFUs at 24 hours relative to the ratio at 0 hours. ***p<0.001 (one-way ANOVA). (E) Growth kinetics of GFP-expressing *L. monocytogenes* (*Lm*-GFP) cocultured with *S. aureus* expressing wild-type S8-Ntox35, catalytic Ntox35 mutant (S8^WT^-Ntox35^H205A^), or empty vector control. Growth of (*Lm*-GFP) was monitored via GFP fluorescence in a microplate reader. (F) Growth kinetics of GFP-expressing *L. monocytogenes* (*Lm*-GFP) in the presence of concentrated supernatants derived from *S. aureus* expressing wild-type S8-Ntox35, catalytic Ntox35 mutant (S8^WT^-Ntox35^H205A^), or heat-inactivated wild-type S8-Ntox35 (95°C for 10 minutes). *Lm*-GFP growth was monitored via GFP fluorescence in a microplate reader. (G) Survival assay of GFP-expressing *L. monocytogenes* (*Lm*-GFP) in the presence of supernatant derived from S. aureus expressing wild-type S8-Ntox35 or catalytic Ntox35 mutant (S8^WT^-Ntox35^H205A^), as determined by colony forming units (CFU) over time. (D-G) Data are presented as mean ± SEM (n = 3 technical replicates, representative of ≥ 2 independent experiments).

The active site of BECR-family RNases contains two conserved histidine residues located on the N-terminal helix (10). Within the BetaH-Ntox35 protein, these correspond to H199 and H205 (Fig. 1B). AlphaFold 3 structural modeling of the Ntox35–immunity protein co-complex predicts a direct interaction between Asp109 of the immunity protein and His205 of Ntox35, consistent with an active site occlusion mechanism in which the immunity protein blocks binding of the toxin to its substrate (Fig. 1C) (12). These structural features strongly support the hypothesis that Ntox35 functions as an RNase toxin with His205 as an active site residue.

Despite this bioinformatic and structural evidence, Ntox35 had not been experimentally validated as a functional antibacterial effector. Notably, within the S8-PTS locus, both the S8 peptidase and the BetaH-Ntox35 fusion protein are predicted to be secreted, whereas the immunity protein lacks a secretion signal, suggesting cytosolic retainment and a potential for kin selection of strains that do not carry immunity (3, 10, 13, 14). To test the antibacterial activity of this system, we conducted competition assays between an *S. aureus* strain expressing the full S8-Ntox35 locus and an isogenic strain lacking this locus (data not shown). Surprisingly, no competitive advantage was observed under these conditions.

Given the presence of the S8-PTS locus in other Firmicutes, we hypothesized that the system may mediate inter-genus competition. To assess this, we utilized *Listeria monocytogenes* 10403S, which lacks a recognizable S8-PTS locus, constitutively expressing GFP (*Lm-*GFP) to facilitate quantification of growth and viability. In contrast to the *S. aureus* intra-species competition, mixed growth on solid media with the S8-Ntox35–expressing *S. aureus* strain resulted in a marked reduction in GFP signal and *Listeria* viability, indicating effective intoxication (Fig. 1D). Importantly, this inhibitory activity was abolished when active site His205 of Ntox35 was mutated to an alanine (H205A), confirming the requirement of the predicted catalytic residue for toxin function (Fig. 1D). Together, these findings demonstrate that the S8-Ntox35 PTS locus encodes a proteinaceous toxin that mediates interbacterial antagonism and that this activity is dependent on the predicted RNase activity of Ntox35.

The presence of an N-terminal signal peptide on Ntox35 suggests that the toxin is exported via the Sec-dependent co-translational secretion pathway (10, 15). Upon secretion the toxin may be retained at the surface of the producing cell and mediate contact dependent inhibition, or it may be released into the extracellular milieu and function as a diffusible inhibitor. The results presented in Fig. 1D do not discriminate between these possibilities. To determine whether the competitive advantage conferred by the S8-Ntox35 locus was mediated by a diffusible effector, we repeated the co-culture competition assay under contact-independent shaking liquid culture conditions. Again, we observed robust inhibition of *L. monocytogenes* in the presence of *S. aureus* expressing the S8-Ntox35 locus, but not with the catalytically inactive S8^WT^-Ntox35^H205A^ mutant (Fig. 1E). To further verify that the inhibitory activity was due to a secreted proteinaceous factor, we collected and concentrated cell-free supernatant from *S. aureus* cultures harboring the S8-Ntox35 locus. This supernatant retained potent growth-inhibitory activity against *L. monocytogenes* that was dependent on a wild-type Ntox35. This activity was abolished by either S8^WT^-Ntox35^H205A^ or by heat treatment of wild-type Ntox35 supernatants at 95°C, consistent with a heat-labile protein (Fig. 1F). Finally, to determine the bacteriostatic or bactericidal activity of Ntox35, we determined *L. monocytogenes* viability following S8-Ntox35 supernatant treatment. Within two hours of exposure, bacterial cell numbers declined by approximately two orders of magnitude relative to the S8^WT^-Ntox35^H205^ mutant, indicating that Ntox35 exerts a bactericidal effect (Fig. 1G). These results confirm the S8-Ntox35 PTS of *S. aureus* functions as a diffusible proteinaceous toxin and further highlight a potentially underappreciated mechanism of inter-genus competition mediated by polymorphic toxins in Firmicutes.

### Toxin Activation Requires S8 Peptidase Processing

S8 peptidases are a class of serine proteases broadly conserved across all domains of life. Many exhibit endopeptidase activity and participate in diverse biological processes, including nutrient acquisition, immune responses, metabolism, and pathogenesis (10, 16, 17). In this PTS locus the S8 peptidase harbors an N-terminal Sec signal peptide. In contrast to typical PTSs, where peptidase domains are embedded within the toxin itself and function to release the C-terminal effector domain during secretion (10), the S8 peptidase in this system is encoded as a separate open reading frame. Nonetheless, its secretion and predicted structure suggest it may contribute to the activity of the associated BetaH-Ntox35 toxin, potentially through post-translational processing or by facilitating translocation into target cells. Structural modeling predicts a direct interaction between the two proteins: a flexible loop within the BetaH domain, located between the β-sheet and α-helix, is positioned within the active site cleft of the S8 peptidase (Fig. 2A). In this model, the catalytic serine (S205) of the peptidase is oriented to mediate nucleophilic hydrolysis of the loop between residues S146 and A147, a position consistent with cleavage between the BetaH domain and the downstream Ntox35 effector.

**Figure 2.**
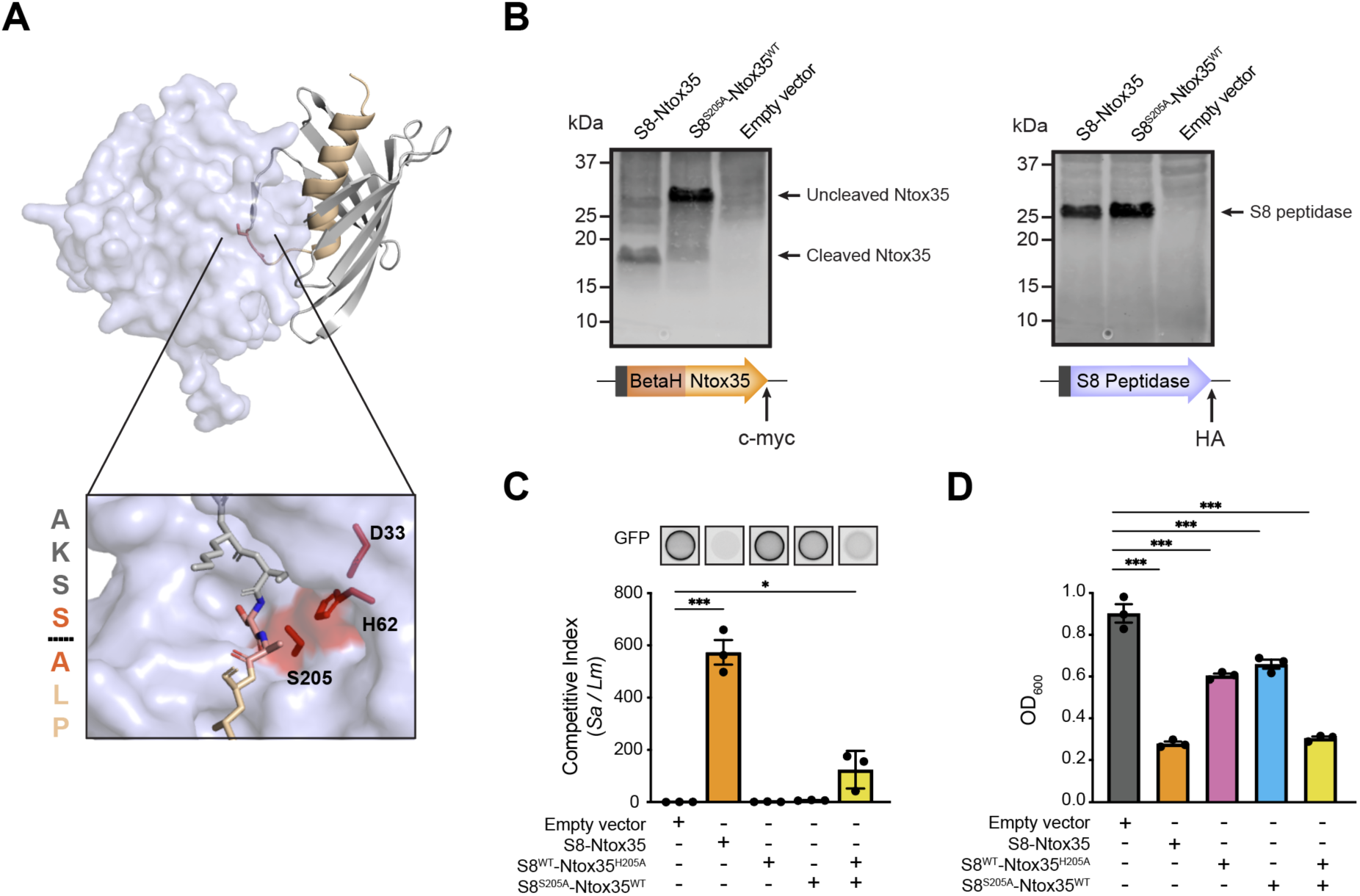
S8-PTS activity requires proteolytic activation of the BetaH-associated toxin by S8 peptidase. (A) AlphaFold 3 modeling of S8 peptidase (blue surface representation) and BetaH domain (grey-tan ribbon structure) interaction. The S8 peptidase active site is placed at a flexible loop separating the β-sheet and α-helix of the BetaH domain. Enlarged inset shows the alignment of S8 S205-H62-D33 catalytic triad in position for proteolytic attack of the amide bond between S146 and A147 of the BetaH domain. Putative cleavage site amino acids listed to the left of the inset with cleavage site indicated by dashed line. (B) Western blot of concentrated supernatants from S. aureus strains expressing wild-type S8-Ntox35, S8^S205A^-Ntox35^WT^, or empty vector control. For all strains, HA and c-myc epitope tags were fused to the C-terminus of the S8 peptidase and Ntox35 proteins, respectively, for western detection by α-myc (Left) and α-HA (Right) antibodies. (C) Contact-dependent competition assay to evaluate the ability of *S. aureus* expressing empty vector, wild-type S8-Ntox35, catalytically dead toxin (S8^WT^-Ntox35^H205A^), catalytically dead peptidase (S8^S205A^-Ntox35^WT^), or a mixture of the two mutant strains to inhibit *L. monocytogenes* expressing GFP (*Lm*-GFP) on solid agar. Representative images show fluorescence of *Lm*-GFP in co-culture with the indicated *S. aureus* strain. Competitive indices were calculated as the ratio of *S. aureus* to *Lm*-GFP CFUs at 24 hours relative to the ratio at 0 hours. (D) Growth kinetics of GFP-expressing *L. monocytogenes* (*Lm*-GFP) in the presence of concentrated supernatants derived from *S. aureus* expressing empty vector, wild-type S8-Ntox35, catalytically dead toxin (S8^WT^-Ntox35^H205A^), catalytically dead peptidase (S8^S205A^-Ntox35^WT^), or a mixture of the two mutant supernatants. Shown is OD_600_ at six hours. (C and D) Data are presented as mean ± SEM (n = 3 technical replicates, representative of ≥ 2 independent experiments). *p<0.05, **p<0.01, and ***p<0.001 (one-way ANOVA).

To experimentally test whether the S8 peptidase mediates cleavage of the BetaH-Ntox35 fusion, we introduced a single amino acid mutation in this predicted catalytic serine to generate a strain with a defective peptidase but wild-type toxin (S8^S205A^-Ntox35^WT^). Additionally, we inserted HA and c-myc epitope tags at the C-terminus of S8 peptidase and Ntox35, respectively, in both the wild-type (S8-Ntox35) and catalytically inactive peptidase (S8^S205A^-Ntox35^WT^) versions of the locus. We reasoned that if the S8 peptidase facilitates toxin release, then wild-type supernatants should contain a cleaved Ntox35 product, whereas supernatants from the S205A mutant should contain the full-length BetaH-Ntox35 fusion. Consistent with this prediction, immunoblot analysis revealed that supernatants from the wild-type S8 strain contained an Ntox35 of approximately 18 kDa, while those from the S8^S205A^ strain contained Ntox35 of approximately 30 kDa (Fig. 2B). These results support a model in which the secreted S8 peptidase cleaves BetaH-Ntox35, releasing the C-terminal toxin domain.

Proteolytic cleavage by the S8 peptidase suggests a role in toxin activation. To determine the requirement of S8 peptidase activity for intoxication, *S. aureus* expressing a functional Ntox35 toxin but harboring the mutant peptidase (S8^S205A^-Ntox35^WT^) (Fig. 2C) or concentrated cell-free supernatants (Fig. 2D) from this strain were applied to susceptible *L. monocytogenes*. In both experiments, mutation of the S8 peptidase active site serine (S205A) abrogated the growth inhibition observed with wild-type Ntox35, mirroring the empty vector or catalytically inactive Ntox35 (Ntox35^H205A^) controls (Fig. 2C-D and Fig. S1). Strikingly, co-culture of strains expressing the inactive Ntox35 toxin (S8^WT^-Ntox35^H205A^) with those harboring the inactive S8 peptidase (S8^S205A^-Ntox35^WT^), as well as incubation with mixed concentrated supernatants derived from those strains, restored the growth-inhibitory phenotype, consistent with trans-complementation between the wild-type toxin and peptidase (Fig. 2C-D and Fig. S1). This result supports a model in which processing of Ntox35 by the active S8 peptidase occurs extracellularly to activate the toxin, reinforcing the hypothesis of a direct functional interaction between the two proteins.

### Bacterial Surface Charge Resists S8-PTS Intoxication

In silico structural analysis of the S8 peptidase–Ntox35 complex predicts that a flexible loop connecting the β-sheet and α-helix of the BetaH domain is positioned near the Asp/Ser/His catalytic triad of the S8 peptidase active site (Fig. 2A). A similar spatial arrangement is predicted in other paired S8-PTS toxins and peptidases (Fig. S2), suggesting a conserved mechanism of interaction and processing. Together with our experimental evidence demonstrating toxin cleavage, these data support a model in which the S8 peptidase mediates proteolytic separation of the N-terminal β-sheet from the α-helix that remains fused to the downstream Ntox35 domain (Fig. 3A). The interface between the α-helix and β-sheet in the intact BetaH domain is stabilized by hydrophobic interactions, while the surface-exposed region of the α-helix and the adjoining loop contain multiple basic residues. In contrast, the β-sheet possesses a strongly anionic surface (10). Based on this, cleavage is predicted to release a cationic and hydrophobic N-terminal extension on the Ntox35 toxin, characteristics reminiscent of cationic antimicrobial peptides (CAMPs), a class of natural antibiotics known to bind and disrupt bacterial membranes (18). To further examine this, we analyzed the hydrophobicity and isoelectric point (pI) of the BetaH domain and its predicted cleavage products across 44 representative BetaH-toxin fusion proteins. Consistently, cleavage resulted in an increase in both hydrophobicity and pI, supporting the notion that S8 processing converts the BetaH domain into a CAMP-like effector (Fig. 3B and C).

**Figure 3.**
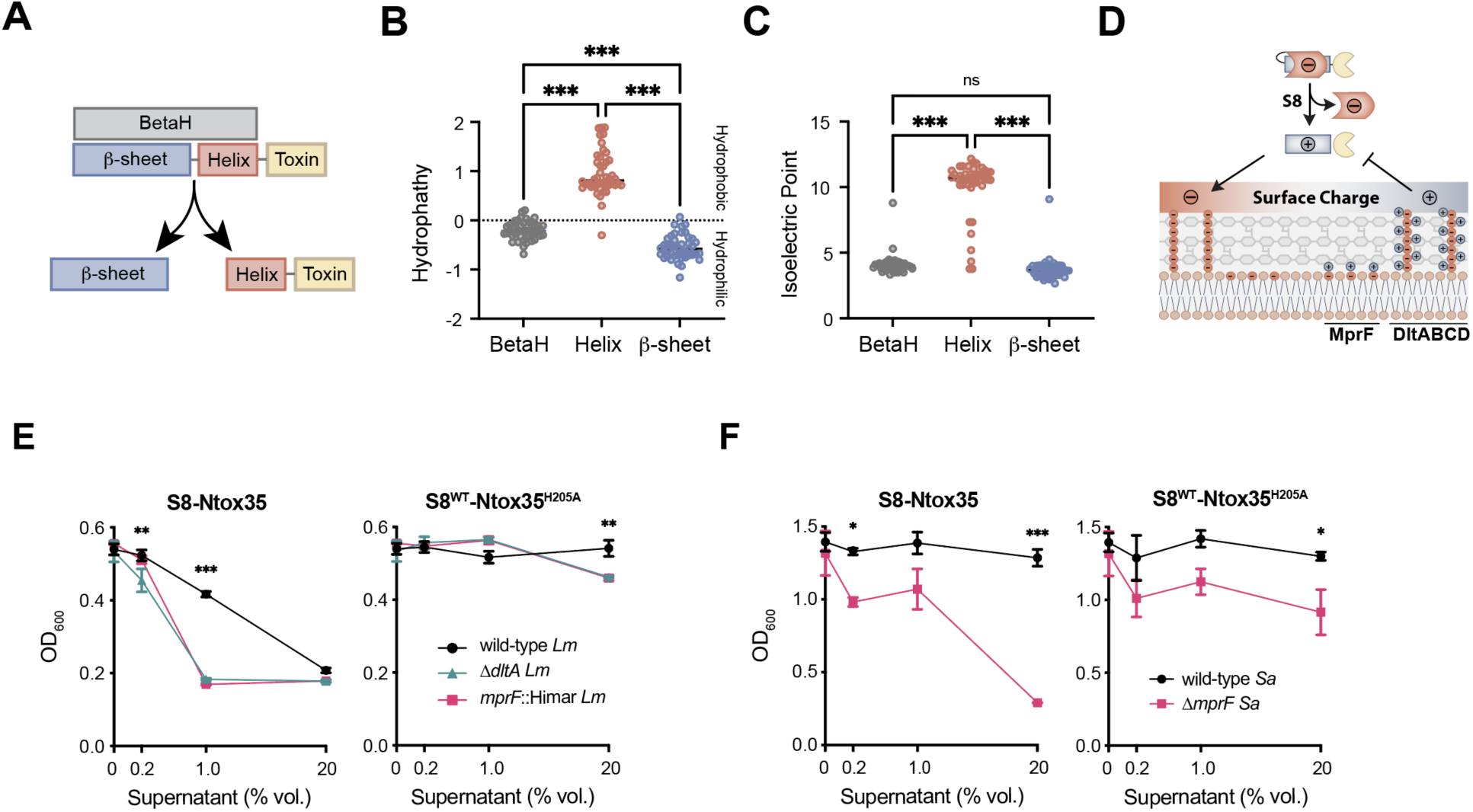
Surface charge modulates toxin susceptibility. (A) Cleavage of the S146-A147 bond would liberate the β-sheet (blue) from the C-terminal α-helix tail (red) and toxin effector (tan) (B) Hydropathy and (C) Isoelectric point determinations of a representative range of forty-four intact BetaH domains (grey), the S8 liberated N-terminal β-sheet (blue), and the C-terminal α-helix tail (red) released with the toxin effector. (D) Cartoon depiction of cell surface charge repulsion of cationic antimicrobial peptides (CAMP), and putative S8 polymorphic toxins. Cell surface charge modifications mediated by lysination of phosphatidylglycerol by MprF or D-alanylation of the cell wall teichoic acid polymers by DltABCD. (E and F) Growth kinetics of wild-type, an *mprF* transposon mutant, and a Δ*dltA* strain of L. *monocytogenes* in the presence of concentrated supernatants derived from *S. aureus* expressing wild-type S8-Ntox35 or S8^WT^-Ntox35^H205A^, at various dilutions. Shown is the OD_600_ at six hours. (G and H) Growth kinetics of wild-type and *ΔmprF* strains of S. aureus in the presence of concentrated supernatants derived from S. aureus expressing wild-type S8-Ntox35 and S8^WT^-Ntox35^H205A^ at various dilutions. Shown is the OD_600_ at six hours. (E-H) Data are presented as mean ± SEM (n = 3 technical replicates, representative of ≥ 2 independent experiments). *p<0.05, **p<0.01, and ***p<0.001 (Two-way ANOVA)

These findings led us to hypothesize that processed Ntox35 may target bacterial cells in a manner analogous to cationic antimicrobial peptides, and that canonical CAMP resistance mechanisms could modulate susceptibility to Ntox35. In Firmicutes, CAMP resistance is commonly mediated by cell surface charge modifications that generate a cationic barrier, repelling positively charged antimicrobials from accessing and disrupting the bacterial membrane. Two well-characterized pathways contribute to this resistance: MprF, which modifies phosphatidylglycerol with lysine (lysyl-PG), and DltABCD, which mediates D-alanylation of wall teichoic acids. Both modifications increase the overall positive surface charge, thereby diminishing CAMP binding and membrane disruption (Fig. 3D) (18–20).

To test whether these pathways influence susceptibility to Ntox35, we challenged wild-type *L. monocytogenes*, a Δ*dltA* mutant (21), and an *mprF* transposon mutant (22) with subinhibitory concentrations of S8-Ntox35–containing supernatant. Both mutants exhibited increased sensitivity to Ntox35-mediated growth inhibition relative to wild-type, with growth of the mutants restricted at 20-fold lower concentrations of toxin containing supernatant than wild-type *L. monocytogenes* (Fig. 3E and Fig. S3). Similarly, *S. aureus*, which was initially resistant to S8-Ntox35 intoxication, became susceptible upon deletion of *mprF*, indicating that surface charge modification protects against Ntox35 activity in this organism even in the absence of a cognate immunity protein (Fig. 3F and Fig. S4). Notably, disruption of these charge-modifying genes (*dltA* and *mprF*) also sensitized both *L. monocytogenes* and *S. aureus* to supernatants containing the catalytically inactive S8^WT^-Ntox35^H205A^ mutant, suggesting that Ntox35 may retain residual, RNase-independent toxicity (Fig. 3E-3F and Fig. S3-S4). These observations support a model in which processed BetaH-Ntox35 functions similarly to CAMPs, whereby target cell susceptibility is influenced by electrostatic interactions with the anionic plasma membrane. Resistance is mediated through charge repulsion, preventing toxin recruitment. Furthermore, our data suggest that residual membrane-disrupting activity, possibly mediated by the cationic, hydrophobic α-helix liberated from the BetaH domain by S8 peptidase cleavage, may contribute to the low-level toxicity of catalytically inactive Ntox35 variants.

### Intoxication by S8-Ntox35 requires the ABC transporter AnrAB

To further investigate the determinants of resistance and susceptibility to Ntox35, we performed an experimental evolution analysis using a sensitive *L. monocytogenes* strain. Specifically, *Lm*-GFP was serially passaged in the presence of subinhibitory concentrations of active Ntox35-containing supernatant. A resistant population emerged after three passages under selective pressure (Fig. 4A) and whole-genome sequencing of nine randomly selected clones from this evolved population was performed. Several changes were observed among the strains (Table S1), however, each strain had at least one missense mutation in at least one component of the VirRS-AnrAB system. VirRS comprise a two-component regulatory system in which VirS is a sensor kinase that phosphorylates the response regulator VirR to impact downstream gene expression. Collectively VirRS mediates resistance to diverse cell envelope damaging antimicrobials (23–25). Downstream regulatory targets of VirRS include MprF and DltABCD, both of which we previously showed to contribute to resistance against S8-Ntox35 (Fig. 3E-F), as well as the ABC-transporter AnrAB, which mediates resistance to the polypeptide antibiotic bacitracin (24, 26–28). Among the evolved strains with resistance to Ntox35 intoxication, one contained a mutation in VirS, one in VirR, and the remaining seven contained one of three missense mutations within the AnrA ATPase component (Fig. 4B).

**Figure 4.**
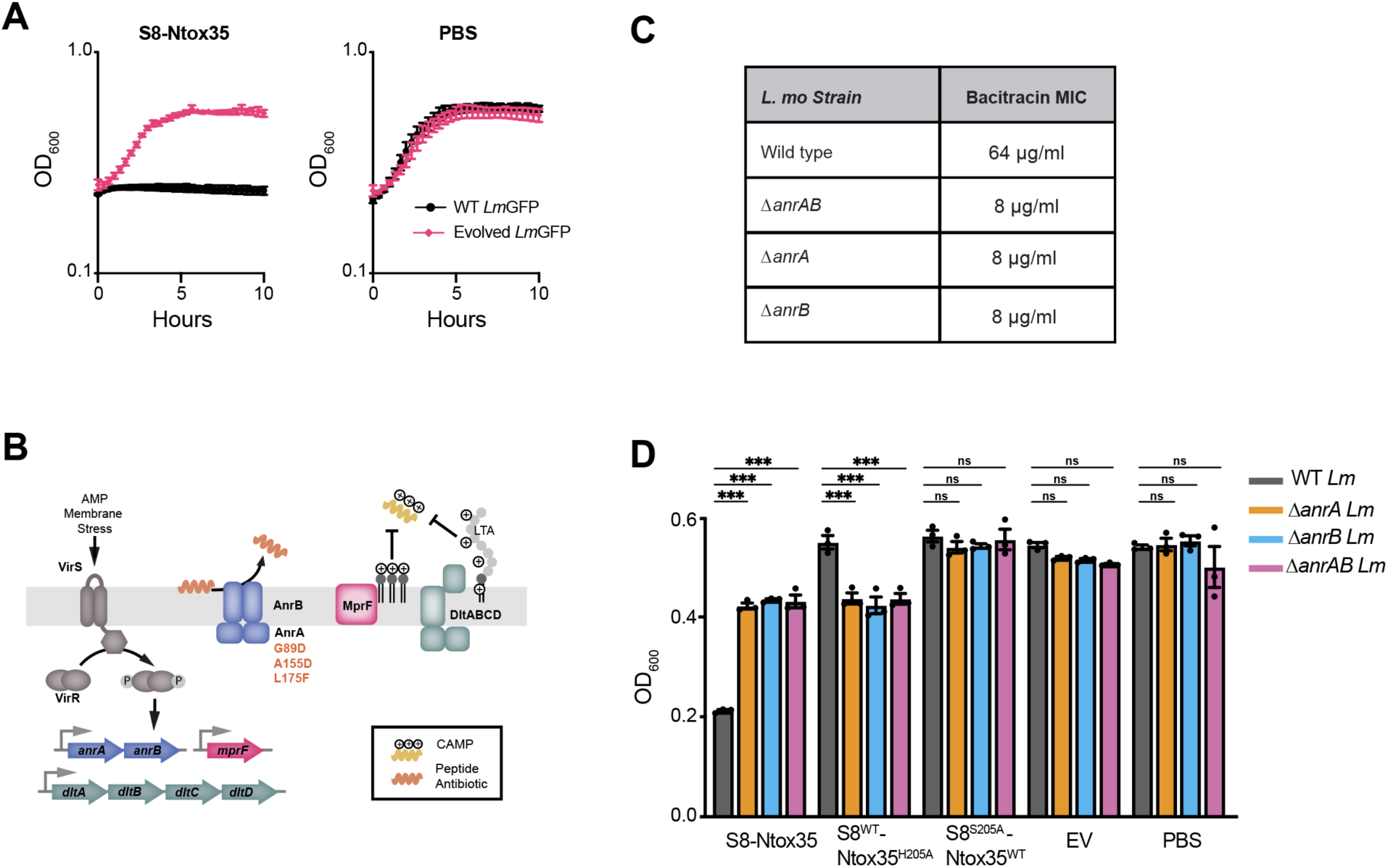
The AnrAB transporter is required for S8-Ntox35 intoxication. (A) Growth kinetics of sensitive GFP-expressing *L. monocytogenes* (*Lm*-GFP) and experimentally evolved *Lm*-GFP, exposed to concentrated supernatant from *S. aureus* expressing wild-type S8-Ntox35 (Left) or PBS (vehicle control) (Right). Data are presented as mean ± SEM (n = 3 technical replicates). (B) Diagram of a cationic antimicrobial peptide detoxification and resistance system within *L. monocytogenes*. Missense mutations in *anrA* from resistant GFP-expressing *L. monocytogenes* are indicated in orange. (C) Bacitracin MIC for wild-type, ΔanrA, Δ*anrB*, and Δ*anrAB L. monocytogenes* strains, determined by broth microdilution (D) Wild-type, Δ*anrA*, Δ*anrB*, and Δ*anrAB L. monocytogenes* strains were exposed to concentrated supernatant from *S. aureus* expressing wild-type S8-Ntox35, the catalytic mutant (S8^WT^-Ntox35^H205A^), empty vector, or PBS (vehicle control). Growth of bacteria was monitored via OD_600_ in a microplate reader, shown is OD_600_ at six hours. Data are presented as mean ± SEM (n = 3 technical replicates, representative of ≥ 2 independent experiments). *p<0.05, **p<0.01, and ***p<0.001 (two-way ANOVA).

Given the established role of the VirRS-AnrAB module in antimicrobial peptide resistance (21–23), we hypothesized that disrupting AnrAB would sensitize *L. monocytogenes* to Ntox35. To test this, we generated complete in-frame deletions of *anrA* and *anrB*, as well as a double knockout strain lacking both genes. To validate the disruption of antimicrobial resistance, we assessed the minimum inhibitory concentration (MIC) of bacitracin using broth microdilution. Consistent with previous reports, deletion of any component of the transporter disrupted bacitracin resistance of *L. monocytogenes*, reducing the MIC from 64 to 8 µg/mL (Fig. 4C) (24–26). Contrary to expectations, however, the combined disruption of AnrAB or either of its individual components conferred increased resistance to S8-Ntox35-containing supernatant (Fig. 4D and Fig. S5). This unexpected result suggested that the AnrAB transporter is essential for Ntox35 susceptibility, potentially functioning in toxin translocation. A role for ABC transporters has previously been reported for translocation of some CDI toxins across the inner membrane of *E. coli*, where disruption of the membrane spanning permease confers toxin resistance (30). Contrary to those findings, however, disruption of the transporter activity is sufficient to protect against Ntox35, as loss of the ATPase AnrA alone phenocopies loss of the permease AnrB. Furthermore, mirroring observations with *dltA* and *mprF* mutants, the *anrAB* mutant strains exhibited increased sensitivity to S8^WT^-Ntox35^H205A^-containing supernatant, despite the toxin’s catalytic inactivation. Strikingly, this susceptibility was lost upon treatment with S8-PTS containing inactive peptidase (S8^S205A^-Ntox35^WT^), consistent with the need to process the BetaH domain (Fig. 4D and Fig. S5). This again suggests that processed Ntox35 possesses residual antimicrobial activity independent of its RNase function, potentially acting extracellularly at the bacterial surface as an antimicrobial peptide. Consequently, AnrAB may attempt to detoxify this effector and inadvertently play a role in facilitating translocation of the catalytic toxin domain of Ntox35.

## Discussion

In this study, we characterized the S8-PTS locus in *Staphylococcus aureus* and demonstrated that it encodes a toxin capable of mediating inter-genus antagonism. We found that the observed bactericidal activity is conferred by a diffusible, heat-labile protein. Furthermore, we showed that toxin activation requires the associated S8 peptidase for extracellular processing, and that bacterial cell surface charge repulsion serves as a mechanism of resistance against S8-Ntox35 intoxication. Finally, we identified a requirement for the ATP-binding cassette (ABC) transporter AnrAB in mediating susceptibility to the associated toxin domain in *L. monocytogenes*.

Beyond its unique function as an inter-genus diffusible antibacterial toxin among Firmicutes, the S8-PTS exhibits several features that distinguish it from other previously characterized systems. First, the use of a separately encoded peptidase for toxin activation contrasts with the more common use of N-terminal endopeptidase domains embedded within polymorphic toxins (3, 10). Moreover, the spontaneous, extracellular nature of toxin processing—occurring independently of target cell contact—raises questions about its functional necessity. We speculate that the β-sheet structure of the BetaH domain cages the cationic hydrophobic α-helical tail to facilitate transit beyond the producing cell envelope and prevent cell surface retention and self-intoxication. Once outside the producing cell, cleavage occurs, exposing a CAMP-like α-helix that facilitates targeting to the membrane of susceptible target cells. As our findings with *dlt* and *mprF* mutants of *L. monocytogenes* and *S. aureus* demonstrate, modulation of target cell surface charge can impact susceptibility via charge repulsion (19, 31).

Experimental evolution of resistance revealed an unexpected role for the ABC protein AnrAB in Ntox35 susceptibility. ABC proteins constitute a large and diverse superfamily of membrane-spanning proteins whose functions, ranging from import to export of numerous molecules, are driven by ATP hydrolysis (32). In some cases, including AnrAB, these systems defend against membrane-targeting agents. AnrAB is co-regulated with *dltABCD* and *mprF* by the VirR response regulator and contributes to resistance against envelope-disrupting antimicrobials, such as bacitracin, by facilitating efflux from the membrane (24–26, 29). Accordingly, disruption of AnrAB was expected to sensitize cells to S8-Ntox35, as seen with other components of the VirR regulon (MprF and DltABCD). Contrary to this expectation, deletion of *anrAB* abrogated the effects of the Ntox35 toxin domain. Although the requirement of the ABC protein for Ntox35 intoxication was unexpected, it is not without precedent. In *Escherichia coli*, membrane components of certain ABC transporters serve as receptors for contact-dependent inhibition (CDI) toxins, while ABC transporters function as receptors of insecticidal Cry toxins produced by *Bacillus thuringiensis.* In CDI systems, ATPase activity is dispensable for intoxication and passage across the membrane may occur passively or by other energetic mechanisms (30, 33, 34). In contrast, our data demonstrate that the AnrA activity is essential for Ntox35-mediated intoxication, an observation consistent with ATPase activity supplying the energy for toxin membrane translocation.

AnrAB is one representative of a conserved group of ABC proteins in Firmicutes that function as resistance determinants against membrane-targeting antimicrobials (28). These systems, often coupled with two-component regulatory systems, form so-called Bce modules— named after the BceRS-BceAB system of *Bacillus subtilis* (24, 25, 35, 36). Bce modules defend against antimicrobial peptides that bind lipid intermediates in cell wall biosynthesis by dislodging and expelling these compounds from the membrane (28, 37). Notably, Bce modules are highly enriched within the Firmicutes phylum, particularly in the Bacillales, Clostridiales, and Lactobacillales orders, where they are found in approximately 97% of sequenced genomes (29). Within Firmicutes, these systems also vary in abundance across species, with some, such as members of Bacillales, encoding multiple Bce-like modules (29, 38).

The co-occurrence of Bce modules among Firmicutes mirrors the phylogenetic distribution of S8-PTS loci, and findings from a parallel study indicate that this toxin family exhibits broad activity across Firmicutes (39). Combined with our data showing that AnrAB is required for S8-PTS intoxication, these findings raise the possibility that Bce modules may play a general role in mediating toxin translocation across susceptible bacteria. However, several lines of evidence complicate this hypothesis. In CDI systems, where toxins similarly require plasma membrane translocation, delivery is typically restricted to recipient cells expressing a conserved membrane receptor (34), suggesting that receptor sequence conservation is a key determinant of translocation specificity. In *Staphylococcus aureus*, the AnrAB homologs VraGF share only ∼25% sequence identity with *L. monocytogenes* AnrAB. Moreover, work by Colautti *et al*. demonstrates S8-PTS-mediated intoxication of *Corynebacterium glutamicum*, an Actinobacterium that lacks a clear homolog of Bce-like ABC transporters. While some orthologous systems have been suggested in *Corynebacteria* (40), the apparent absence of conserved membrane protein sequences challenges the idea that AnrAB-like transporters universally act as direct translocators for S8-PTS toxins.

Despite limited sequence conservation, these transporters may share a functional role in defending the cell envelope—specifically, by protecting undecaprenyl pyrophosphate (UPP), a conserved lipid intermediate required for peptidoglycan synthesis. It is conceivable that S8-PTS effectors exploit this common lipid target and co-opt the activity of Bce modules during attempts to release UPP, thereby entering the cell. This model may explain why *S. aureus*, despite harboring a low-identity AnrAB homolog, becomes susceptible to Ntox35 following deletion of *mprF*. Further investigation into the role of Bce modules in S8-PTS intoxication across Firmicutes beyond *L. monocytogenes* will clarify this from other mechanisms by which AnrAB imparts S8-PTS susceptibility.

Like other PTSs, Ntox35 intoxication ultimately relies on delivery of a C-terminal toxin domain into the target cell cytosol. However, S8 peptidase–mediated processing uniquely liberates a caged, cationic amphipathic α-helix. Our data suggest this peptide serves a dual function: facilitating plasma membrane translocation of the toxin and directly disrupting bacterial growth independent of RNase activity. This later activity is supported by the increased sensitivity of *anrAB* mutants to the catalytically inactive S8^WT^-Ntox35^H205A^ variant but not those with the mutant peptidase S8^S205A^-Ntox35^WT^. In this context, the requirement for AnrAB in Ntox35 susceptibility highlights an evolutionary double-edged sword in which loss of AnrAB confers resistance to S8-PTS toxin domain mediated killing but concurrently increases sensitivity to membrane-targeting antimicrobials, including the membrane-disrupting component of the processed Ntox35 toxin itself.

Bioinformatic studies, such as those by Li *et al.* (10), have revealed the widespread prevalence and diversity of PTSs across both Gram-negative and Gram-positive bacteria. However, experimental characterization in Gram-positive species has thus far remained limited. Although the findings presented here and those addressed in parallel by Colautti *et al*., including biochemical confirmation of Ntox35 RNase activity, mapping of S8 peptidase cleavage sites, and breadth of toxicity, offer important initial insights; a comprehensive understanding of the molecular mechanisms underlying S8-PTS–mediated bacterial antagonism will require continued effort. Beyond future molecular details of the S8-PTS, *in vivo* investigation of the impacts of the S8-PTS on interbacterial competition represents a critical next step toward understanding its broader ecological and clinical significance within the microbiota and polymicrobial infections. We anticipate that this work will provide a foundation for future studies aimed at surveying S8-PTS systems across Firmicutes, characterizing their functional diversity, and uncovering the evolutionary pressures shaping their specificity and distribution.

## Methods

### Quantification and Statistical Analysis

Statistical analyses were performed using GraphPad Prism 10. Details on the statistical tests used, number of replicates, and definition of center and error bars are found in the figure legends. Results with *p*<0.05 are considered statistically significant, *p<0.05, **p<0.01, and ***p<0.001.

### Bacterial Strains and Culture Conditions

All bacterial strains used in this study are summarized in Table S2. Bacteria were cultured on Brain Heart Infusion (BHI) (RPI B11000-5000.0) agar (*L. monocytogenes*, *S. aureus*) or LB Miller agar (Fisher Scientific BP97235) (*L. monocytogenes*, *S. aureus*, *E*. *coli*) at 37°C. Unless otherwise stated, prior to experiments bacteria were grown in BHI broth (*L. monocytogenes*, *S. aureus*) or LB Miller broth overnight at 37°C with shaking. Antibiotics (RPI) were used at the following concentrations: streptomycin, 200 μg/mL (*L. monocytogenes*); chloramphenicol, 10 μg/mL (*E*. *coli*, *S. aureus*); erythromycin, 2 µg/mL (*L. monocytogenes*); and carbenicillin, 100 μg/mL (*E*. *coli*). Plasmids were introduced into *E. coli* by chemical transformation, into *L. monocytogenes* using trans-conjugation, and into *S. aureus* by electroporation.

### Plasmid and Strain Construction

Plasmids and oligonucleotides used in this study are listed in Tables S3 and S4, respectively. Primers and synthetic DNA fragments were obtained from Integrated DNA Technologies. All plasmid constructs were designed using Snap Gene and generated using Gibson Assembly, In-Fusion cloning, or restriction enzyme cloning, and all constructs were confirmed by sequencing. Amino acid substitutions were generated using overlapping site directed mutagenesis primers. Substitutions were confirmed by PCR, Sanger sequencing, and whole plasmid sequencing. In-frame unmarked gene deletions in *L. monocytogenes* were generated via allelic exchange using plasmid pLIM followed by counterselection on 18 mM DL-4-chlorophenylalanine (41) and were confirmed by Sanger sequencing of PCR amplicons spanning the gene disruption. Plasmids were introduced to *L. monocytogenes* via conjugation from E. coli SM10 (42). In-frame unmarked gene deletions in *S. aureus* were generated via allelic exchange using plasmid pIMAY*-Z (43) and inserts were confirmed by PCR and Sanger sequencing. Plasmid DNA was extracted from *E. coli* IM08B and introduced to *S. aureus* by electroporation and maintained by antibiotic selection (44–46). pEPSA5 (47) was used to express S8-Ntox35 locus constructs in *S. aureus*, without xylose induction of the plasmid.

### Preparation of Concentrated Supernatant for S8-Ntox35 Toxicity Assays

*Staphylococcus aureus* strains harboring either pEPSA5::S8-Ntox35-TPR_Imm1 or a corresponding mutant construct were used to produce toxin and peptidase containing supernatant (47). A single colony was inoculated into 100 mL of Brain Heart Infusion (BHI) broth and incubated, without xylose induction of pEPSA5, at 37 °C with shaking for 20 hours. Cultures were then centrifuged at 4,500 xg for 20 minutes to pellet the cells. The resulting supernatant was sterile-filtered through a 0.2 µm vacuum-driven membrane filter (Thermo Scientific 565-0020) to remove any remaining bacterial cells. The clarified supernatant was then concentrated using a 10 kDa molecular weight cutoff Amicon Ultra Centrifugal Filter Unit (Millipore UFC901024) until the volume was reduced to approximately 2.5 mL. To remove small molecules and buffer exchange into phosphate-buffered saline (PBS), the concentrate was processed through a Cytiva PD-10 desalting column (17-0851-01) according to the manufacturer’s instructions and sterile-filtered through a 0.2 µm syringe filter (Sterlitech 1470353) The final product was aliquoted, snap-frozen in liquid nitrogen, and stored at −80 °C until further use.

### Measurement of S8-Ntox35 Susceptibility

To assess susceptibility to S8-Ntox35, cultures of the indicated bacterial strains were grown in the appropriate media overnight at 37 °C with shaking. Stationary phase cultures were adjusted to an optical density of 600 nm (OD₆₀₀) of 0.25 using fresh BHI or the relevant growth medium. For each condition, 80 µL of the diluted culture was dispensed into wells of a sterile 96-well cell culture plate (GenClone 25-104) in triplicate. Subsequently, 20 µL of various dilutions of either concentrated S8-Ntox35 supernatant or control supernatant was added to each well, yielding a total volume of 100 µL per well. Plates were sealed with BreatheEasy® gas-permeable film (Diversified Biotech BEM-1) and incubated at the indicated temperature with continuous shaking in a BioTek Synergy HTX Multi-Mode Microplate Reader (Agilent). Optical density at 600 nm (OD₆₀₀) and GFP-associated relative fluorescence units (RFU) (485/528), when applicable, were recorded every 20 minutes throughout the incubation period.

### Bactericidal Assay

The impacts of S8-Ntox35 on bacterial viability were assessed using a modified version of the measurement of S8-Ntox35 susceptibility. Overnight bacterial cultures were diluted to an OD₆₀₀ of 0.25 in fresh BHI. For each condition, 80 µL of the diluted culture was mixed with 20 µL of concentrated S8-Ntox35 or control supernatant in a 96-well microtiter plate. Plates were incubated at the indicated temperature with shaking in a standard benchtop incubator. At 1- and 2-hours post-initiation, 10 µL samples were taken from each well, serially diluted in PBS, and plated on appropriate selective media for enumeration of colony-forming units (CFUs). Input CFUs (0-hour timepoint) were similarly determined by plating samples collected immediately after supernatant addition. Plates were incubated overnight at 37 °C before counting.

### Contact Competition

To assess interspecies contact-dependent competition, *Listeria monocytogenes* 10403S expressing GFP (*Lm*-GFP) and *Staphylococcus aureus* were grown overnight in Brain Heart Infusion (BHI) broth at 37 °C with agitation. The following day, cultures were normalized to an optical density at 600 nm (OD₆₀₀) of 0.1 in fresh BHI. Equal volumes of each normalized culture were mixed and 5 µL of the mixture was spotted onto LB agar plates. Plates were incubated overnight at 37 °C. The following day, colonies were imaged using an iBright FL1500 Imaging System (Thermo Fisher Scientific) to visualize GFP fluorescence and assess the presence of *Lm*-GFP. Colonies were excised from the agar using a sterile scalpel and resuspended in phosphate-buffered saline (PBS). A sample from the suspension was serially diluted in PBS and plated on on appropriate selective media for enumeration of colony-forming units (CFUs). The competitive index (CI) was calculated as the ratio of *S. aureus* to *L. monocytogenes* CFUs at 24 hours divided by the ratio of *S. aureus* to *L. monocytogenes* CFUs in the input mix.

### Suspension Competition

To evaluate bacterial competition in liquid culture, *Listeria monocytogenes* constitutively expressing (*Lm*-GFP) and *Staphylococcus aureus* were grown overnight in Brain Heart Infusion (BHI) broth at 37 °C with agitation. Overnight cultures were diluted to an OD₆₀₀ of 0.1 in fresh BHI, and equal volumes of each strain were combined in wells of a sterile 96-well cell culture plate (GenClone 25-104). Each well contained a final volume of 100 µL. The plate was sealed with BreathEasy® gas-permeable sealing film (Diversified Biotech BEM-1) and incubated at 37 °C with continuous shaking in a BioTek Synergy HTX Multi-Mode Microplate Reader (Agilent) for 20 hours. Optical density at 600 nm (OD₆₀₀) and GFP-associated relative fluorescence units (RFU) (485/528) were measured at regular intervals using the absorbance and fluorescence detection settings, respectively.

### Experimental Evolution

To select for resistance to the S8-Ntox35 effector, a single colony of Ntox35-sensitive *Listeria monocytogenes* constitutively expressing GFP (*Lm*-GFP) was inoculated into Brain Heart Infusion (BHI) broth and grown overnight at 37 °C with shaking. The following day, the culture was back-diluted 1:100 into 1 mL of fresh BHI, and 100 µL of concentrated S8-Ntox35– containing supernatant was added. The culture was incubated overnight at 37 °C with shaking. This procedure was repeated for three consecutive overnight passages. After the final passage, the evolved culture was assessed for Ntox35 susceptibility as described for the suspension competition. To isolate individual clones from the evolved population, the culture was serially diluted in phosphate-buffered saline (PBS) and plated on BHI agar to obtain single colonies. Ten colonies were randomly selected for whole-genome sequencing, of which nine were successfully completed. Library preparation was performed using the Illumina DNA Prep kit, and samples were sequenced on the iSeq 100 Sequencing System. FASTQ files obtained from sequencing were analyzed using the Variation Analysis tool available on the BV-BRC Resources platform (48, 49). Sequencing results are summarized in Table S1.

### Protein SDS-PAGE and Western blotting

To detect tagged proteins in the concentrated supernatant, samples were mixed with 4× Laemmli buffer containing β-mercaptoethanol (BME) to a final 1× concentration, then boiled at 95 °C for 10 minutes. Samples were briefly centrifuged before being loaded onto Bio-Rad TGX Stain-Free 4–20% Mini-PROTEAN (4569036) gels. Precision Plus Protein™ All Blue Prestained Protein Standards were loaded alongside. Electrophoresis was performed at 100 V for 15 minutes, followed by 150 V for 35 minutes. Proteins were transferred onto nitrocellulose using a Trans-Blot Turbo Nitrocellulose Transfer Pack (1704158) and the Bio-Rad Trans-Blot Turbo Transfer System. Total protein was visualized using the LI-COR Revert Total Protein Stain (926-11010) according to the manufacturer’s protocol and imaged on the LI-COR Odyssey Fc Imaging System. Membranes were then blocked for 1 hour at room temperature with gentle shaking in LI-COR Intercept TBS Blocking Buffer (927-60001) supplemented with human serum (Complement Technology NHS, 1:500). Membranes were incubated overnight at 4 °C with rabbit polyclonal anti-Myc antibody (Proteintech 16286-1-AP, 1:5000) and mouse monoclonal anti-HA antibody (Abcam ab18181, 1:5000) diluted in blocking buffer. Following incubation, membranes were washed three times for 7 minutes each with TBS-T (Tris-buffered saline, 0.1% Tween-20), then incubated with IRDye 800CW goat anti-rabbit IgG and IRDye 680CW goat anti-mouse IgG secondary antibody (LI-COR 926-32211 and 926-68070, 1:15,000) for 1 hour at room temperature. After three additional washes with TBS-T and a final rinse in TBS, blots were imaged using the LI-COR Odyssey CLx Imaging System.

### *In silico* analysis of S8-PTS components

Structural predictions of individual proteins and protein-protein complexes were conducted using AlphaFold 3 (50). Structural similarity search was performed using the Dali server (51). Protein sequences used for analysis of interactions between BetaH-toxins and cognate immunity and S8 peptidases are provided in Table S5. The sequence of forty four distinct BetaH-toxins discussed by Li et al. (10) were used for the determination of hydropathy and isoelectric point of the intact BetaH domain and the two resulting fragments generated by S8 protease processing. The analyzed sequences are provided in Table S6. Isoelectric points were determined using IPC (52) and the grand average of hydropathy was determined using the online tool (www.gravy-calculator.de).

## Acknowledgements

We would like to thank Stephen Salipante and Lucas Hoffman for providing *S. aureus* strains containing the S8-Ntox35 PTS, Daniel Portnoy for the *mprF*::Tn and Δ*dltA L. monocytogenes* strains, Joseph Mougous and Savannah Farrell for DNA library preparation and whole genome sequencing of suppressor mutations, Pete Lauer for GFP expressing *L. monocytogenes,* John Whitney for constructive feedback on the manuscript, and the Reniere and Woodward labs for helpful discussions. This work was supported by NIH/NIAID grants R01AI116669 and R01AI139071.

---

**Figure S1:**
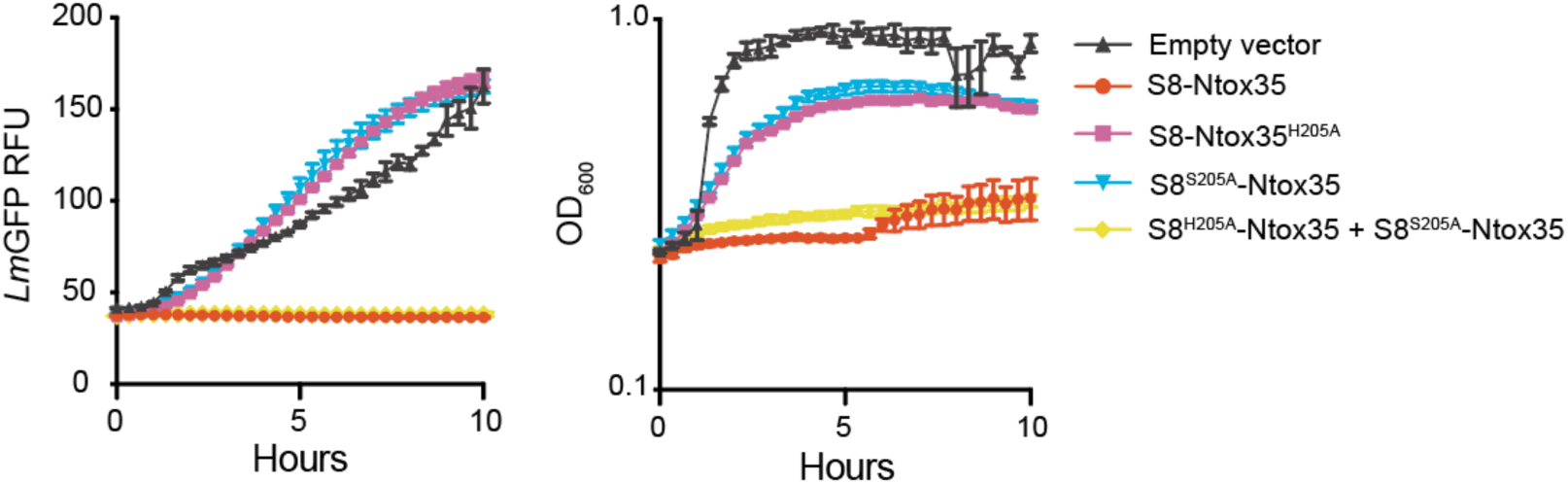
Full growth curves for data presented in Figures 2C-D. Ten-hour broth growth curves of *Lm-GFP* grown in the presence of indicated *S. aureus* supernatants. GFP signal and optical density are depicted.

**Figure S2:**
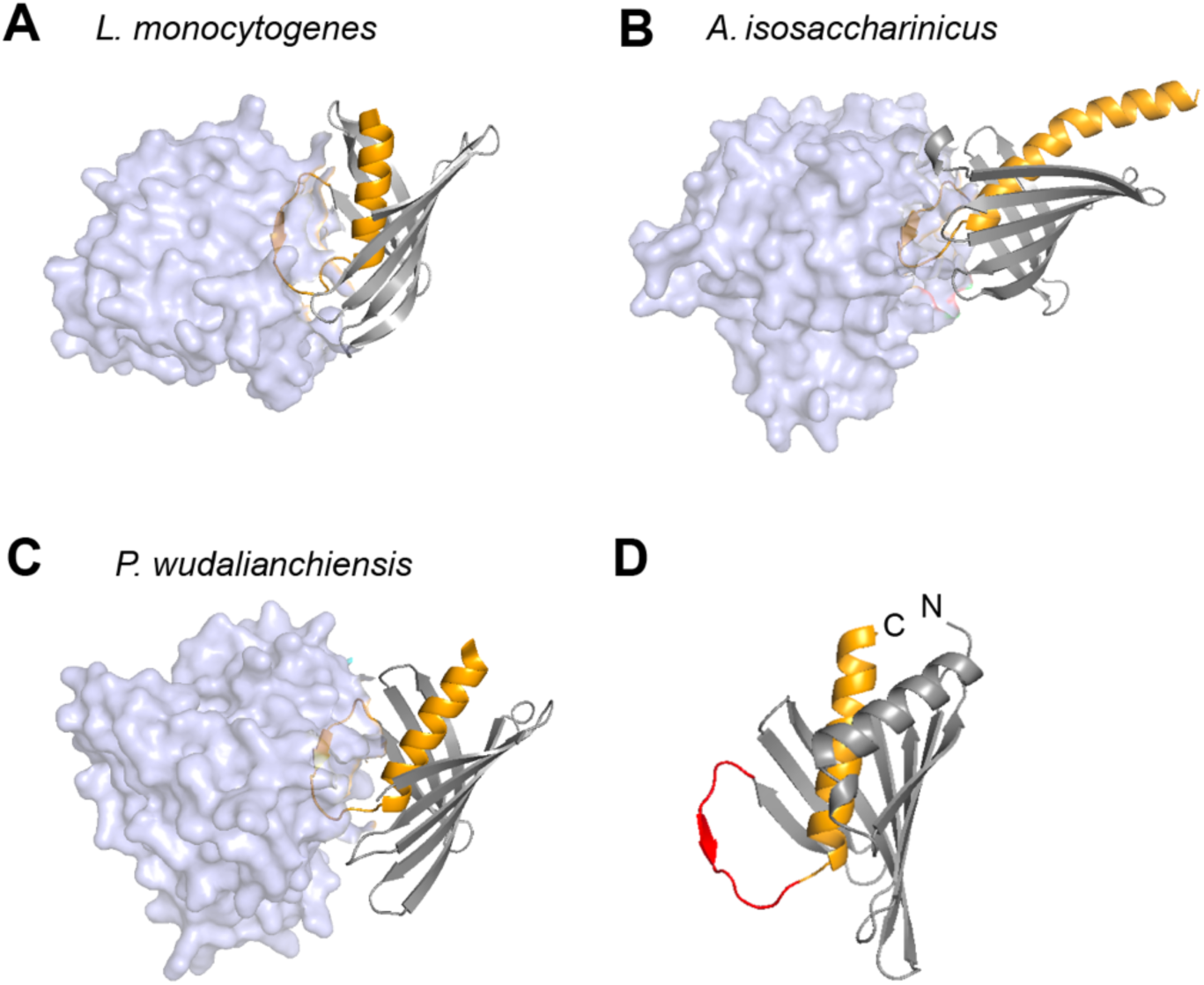
Structural prediction of peptidase-toxin complexes. The S8 peptidase and cognate toxin of the S8-PTS locus found in the indicated bacteria were subjected to AlphaFold 3 modeling. (A-C) The BetaH domain comprised of a β-sheet (gray cartoon) and the caged α -helix (orange) in complex with S8 peptidase (transparent blue surface) are depicted. (D) The flexible loop connecting the β -sheet and α -helix of the BetaH domain reaching into the cleft of the S8 peptidase active site for the *Pseudobacillus wudalianchiensis* toxin is depicted in red.

**Figure S3:**
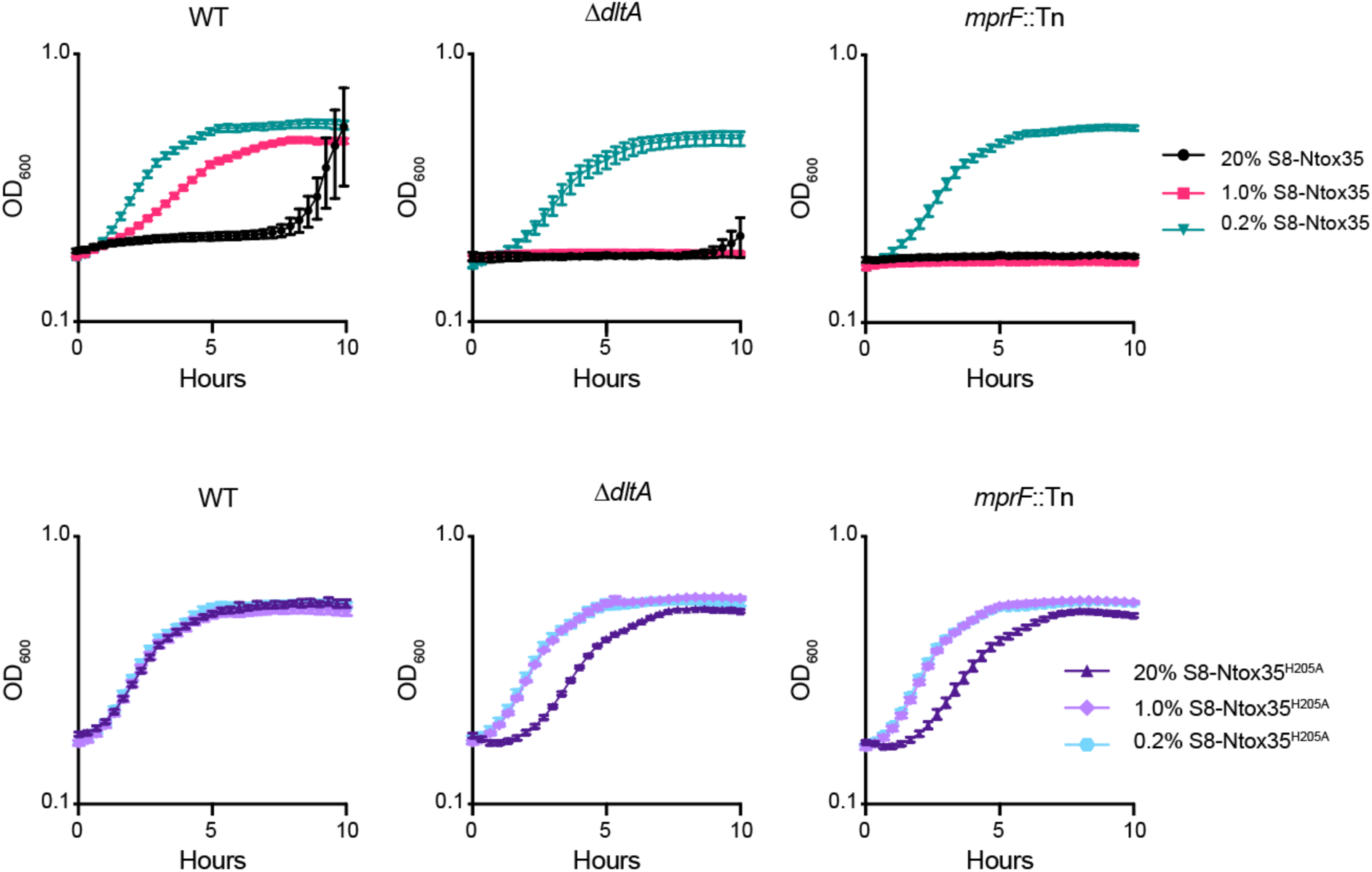
Full growth curves for data presented in Figure 3E. Ten-hour broth growth curves of indicated *L. monocytogenes* strains grown in the presence of various concentrations of the indicated *S. aureus* supernatants. Dilutions represent the % total volume of supernatant for total culture volume. Values are OD_600_ measured using a BioTek plate reader every 20 minutes, Data are presented as mean ± SEM (n = 3 technical replicates, representative of ≥ 2 independent experiments).

**Figure S4:**
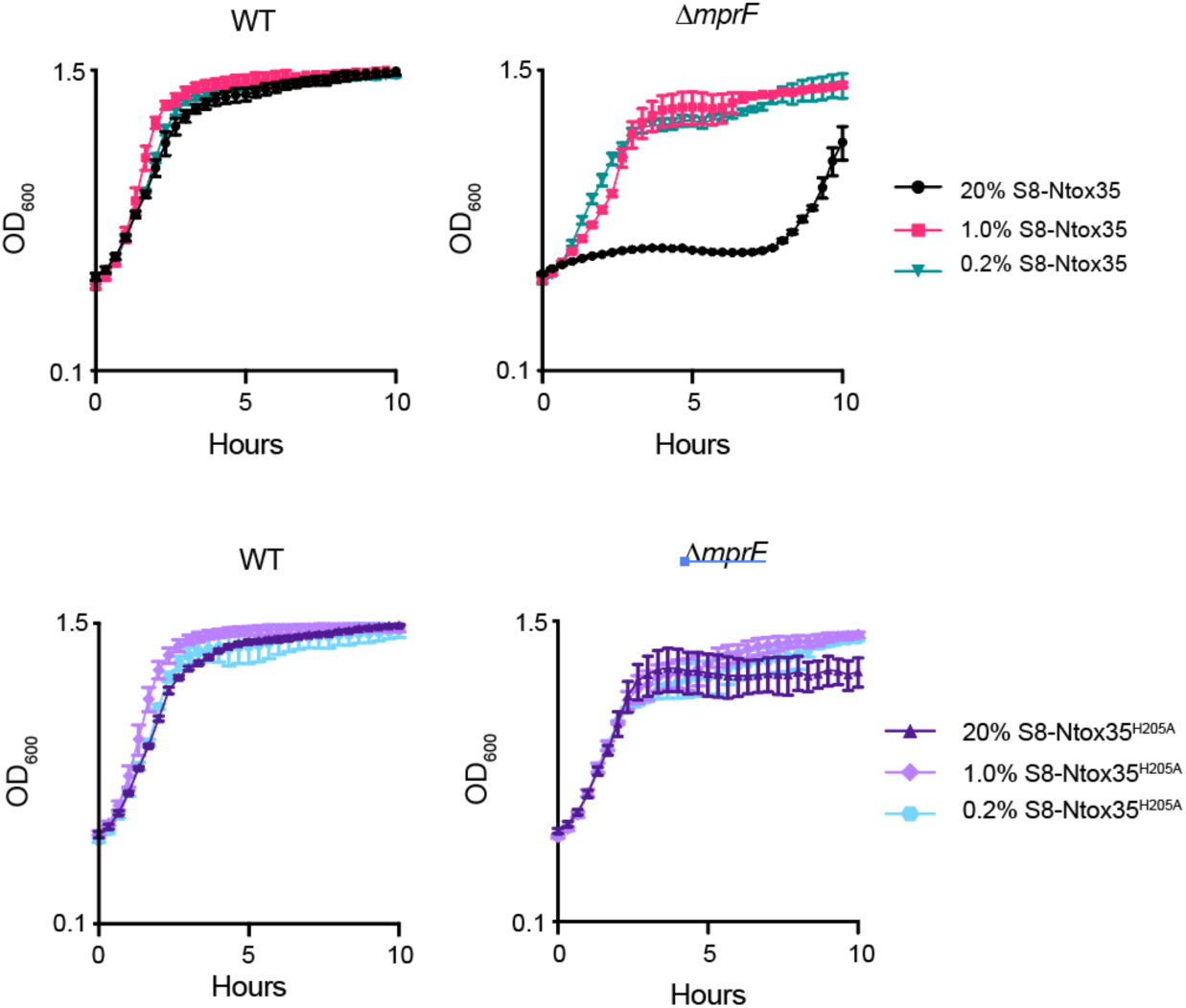
Full growth curves for *S. aureus* data presented in Figure 3F. Ten-hour broth growth curves of indicated *S. aureus* strains grown in the presence of various concentrations of the indicated *S. aureus* supernatants. Dilutions represent the % total volume of supernatant for total culture volume. Values are OD_600_ measured using a BioTek plate reader every 20 minutes, Data are presented as mean ± SEM (n = 3 technical replicates, representative of ≥ 2 independent experiments).

**Figure S5:**
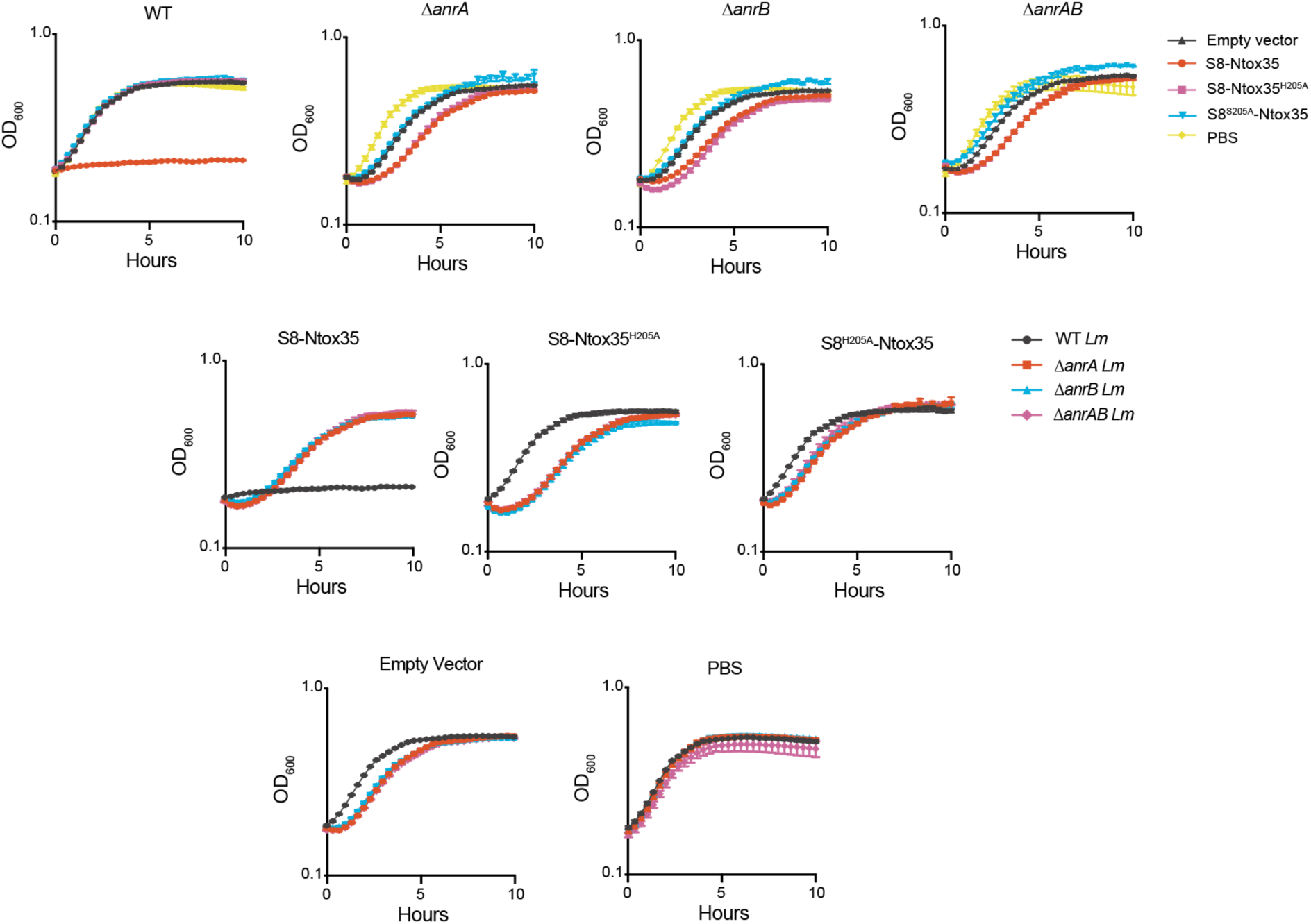
Full growth curves for data presented in Figure 4C. Ten-hour broth growth curves of indicated *L. monocytogenes* strains grown in the presence of the indicated *S. aureus* supernatants. Curves are plotted (i) as individual strains with each indicated treatment and (ii) as each individual treatment with the four strains under investigation. Values are OD_600_ measured using a BioTek plate reader every 20 minutes, Data are presented as mean ± SEM (n = 3 technical replicates, representative of ≥ 2 independent experiments).

## REFERENCES

1. E. T. Granato, T. A. Meiller-Legrand, K. R. Foster, The Evolution and Ecology of Bacterial Warfare. Current Biology 29, R521–R537 (2019).

2. S. K. Aoki, et al., A widespread family of polymorphic contact-dependent toxin delivery systems in bacteria. Nature 468, 439–442 (2010).

3. D. Zhang, R. F. De Souza, V. Anantharaman, L. M. Iyer, L. Aravind, Polymorphic toxin systems: Comprehensive characterization of trafficking modes, processing, mechanisms of action, immunity and ecology using comparative genomics. Biol Direct 7, 18 (2012).

4. S. K. Aoki, et al., Contact-Dependent Inhibition of Growth in *Escherichia coli*. Science 309, 1245–1248 (2005).

5. J. C. Whitney, et al., A broadly distributed toxin family mediates contact-dependent antagonism between gram-positive bacteria. eLife 6, e26938 (2017).

6. R. D. Hood, et al., A Type VI Secretion System of Pseudomonas aeruginosa Targets a Toxin to Bacteria. Cell Host Microbe 7, 25–37 (2010).

7. E. Cascales, et al., Colicin Biology. Microbiol Mol Biol Rev 71, 158–229 (2007).

8. Q. Zhao, et al., Streptomyces umbrella toxin particles block hyphal growth of competing species. Nature 629, 165–173 (2024).

9. A. Jamet, X. Nassif, New Players in the Toxin Field: Polymorphic Toxin Systems in Bacteria. mBio 6, e00285–15 (2015).

10. H. Li, Y. Tan, D. Zhang, Genomic discovery and structural dissection of a novel type of polymorphic toxin system in gram-positive bacteria. Computational and Structural Biotechnology Journal 20, 4517–4531 (2022).

11. B. E. Heaton, J. Herrou, A. E. Blackwell, V. H. Wysocki, S. Crosson, Molecular Structure and Function of the Novel BrnT/BrnA Toxin-Antitoxin System of Brucella abortus. Journal of Biological Chemistry 287, 12098–12110 (2012).

12. Z. C. Ruhe, D. A. Low, C. S. Hayes, Polymorphic Toxins and Their Immunity Proteins: Diversity, Evolution, and Mechanisms of Delivery. Annu. Rev. Microbiol. 74, 497–520 (2020).

13. S. Kaundal, A. Deep, G. Kaur, K. G. Thakur, Molecular and Biochemical Characterization of YeeF/YezG, a Polymorphic Toxin-Immunity Protein Pair From Bacillus subtilis. Front. Microbiol. 11, 95 (2020).

14. D. Zhang, L. M. Iyer, L. Aravind, A novel immunity system for bacterial nucleic acid degrading toxins and its recruitment in various eukaryotic and DNA viral systems. Nucleic Acids Research 39, 4532–4552 (2011).

15. O. Schneewind, D. Missiakas, Sec-secretion and sortase-mediated anchoring of proteins in Gram-positive bacteria. Biochimica et Biophysica Acta (BBA) - Molecular Cell Research 1843, 1687–1697 (2014).

16. F. Falkenberg, M. Bott, J. Bongaerts, P. Siegert, Phylogenetic survey of the subtilase family and a data-mining-based search for new subtilisins from Bacillaceae. Front. Microbiol. 13 (2022).

17. N. D. Rawlings, A. J. Barrett, A. Bateman, MEROPS: the peptidase database. Nucleic Acids Research 38, D227–D233 (2010).

18. S. Omardien, S. Brul, S. A. J. Zaat, Antimicrobial Activity of Cationic Antimicrobial Peptides against Gram-Positives: Current Progress Made in Understanding the Mode of Action and the Response of Bacteria. Front. Cell Dev. Biol. 4 (2016).

19. K. Thedieck, et al., The MprF protein is required for lysinylation of phospholipids in listerial membranes and confers resistance to cationic antimicrobial peptides (CAMPs) on *Listeria monocytogenes*. Molecular Microbiology 62, 1325–1339 (2006).

20. F. C. Neuhaus, J. Baddiley, A Continuum of Anionic Charge: Structures and Functions ofD-Alanyl-Teichoic Acids in Gram-Positive Bacteria. Microbiol Mol Biol Rev 67, 686–723 (2003).

21. T. P. Burke, et al., Listeria monocytogenes Is Resistant to Lysozyme through the Regulation, Not the Acquisition, of Cell Wall-Modifying Enzymes. J Bacteriol 196, 3756– 3767 (2014).

22. J. Zemansky, et al., Development of a*mariner*-Based Transposon and Identification of*Listeria monocytogenes*Determinants, Including the Peptidyl-Prolyl Isomerase PrsA2, That Contribute to Its Hemolytic Phenotype. J Bacteriol 191, 3950–3964 (2009).

23. J. Kang, M. Wiedmann, K. J. Boor, T. M. Bergholz, VirR-Mediated Resistance of Listeria monocytogenes against Food Antimicrobials and Cross-Protection Induced by Exposure to Organic Acid Salts. Appl Environ Microbiol 81, 4553–4562 (2015).

24. X. Jiang, et al., The VirAB-VirSR-AnrAB Multicomponent System Is Involved in Resistance of Listeria monocytogenes EGD-e to Cephalosporins, Bacitracin, Nisin, Benzalkonium Chloride, and Ethidium Bromide. Appl Environ Microbiol 85 (2019).

25. D. Grubaugh, et al., The VirAB ABC Transporter Is Required for VirR Regulation of Listeria monocytogenes Virulence and Resistance to Nisin. Infect Immun 86 (2018).

26. B. Collins, N. Curtis, P. D. Cotter, C. Hill, R. P. Ross, The ABC Transporter AnrAB Contributes to the Innate Resistance of *Listeria monocytogenes* to Nisin, Bacitracin, and Various β-Lactam Antibiotics. Antimicrob Agents Chemother 54, 4416– 4423 (2010).

27. P. Mandin, et al., VirR, a response regulator critical for *Listeria monocytogenes* virulence. Molecular Microbiology 57, 1367–1380 (2005).

28. J. Rismondo, L. M. Schulz, Not Just Transporters: Alternative Functions of ABC Transporters in Bacillus subtilis and Listeria monocytogenes. Microorganisms 9, 163 (2021).

29. S. Dintner, et al., Coevolution of ABC Transporters and Two-Component Regulatory Systems as Resistance Modules against Antimicrobial Peptides in Firmicutes Bacteria. J Bacteriol 193, 3851–3862 (2011).

30. J. L. E. Willett, G. C. Gucinski, J. P. Fatherree, D. A. Low, C. S. Hayes, Contact-dependent growth inhibition toxins exploit multiple independent cell-entry pathways. Proc. Natl. Acad. Sci. U.S.A. 112, 11341–11346 (2015).

31. E. Abachin, et al., Formation of D-alanyl-lipoteichoic acid is required for adhesion and virulence of *Listeria monocytogenes*. Molecular Microbiology 43, 1–14 (2002).

32. M. J. Fath, R. Kolter, ABC transporters: bacterial exporters. Microbiol Rev 57, 995–1017 (1993).

33. R. Sato, S. Adegawa, X. Li, S. Tanaka, H. Endo, Function and Role of ATP-Binding Cassette Transporters as Receptors for 3D-Cry Toxins. Toxins 11, 124 (2019).

34. T. Myers-Morales, M. M. S. Sim, T. J. DuCote, E. C. Garcia, *Burkholderia multivorans* requires species-specific GltJK for entry of a contact-dependent growth inhibition system protein. Molecular Microbiology 116, 957–973 (2021).

35. A. Staroń, D. E. Finkeisen, T. Mascher, Peptide Antibiotic Sensing and Detoxification Modules of *Bacillus subtilis*. Antimicrob Agents Chemother 55, 515–525 (2011).

36. S. Dintner, R. Heermann, C. Fang, K. Jung, S. Gebhard, A Sensory Complex Consisting of an ATP-binding Cassette Transporter and a Two-component Regulatory System Controls Bacitracin Resistance in Bacillus subtilis. Journal of Biological Chemistry 289, 27899– 27910 (2014).

37. N. L. George, A. L. Schilmiller, B. J. Orlando, Conformational snapshots of the bacitracin sensing and resistance transporter BceAB. Proc. Natl. Acad. Sci. U.S.A. 119 (2022).

38. S. Gebhard, T. Mascher, Antimicrobial peptide sensing and detoxification modules: unravelling the regulatory circuitry of *Staphylococcus aureus*. Molecular Microbiology 81, 581–587 (2011).

39. J. Colautti, S. R. Garrett, J. C. Whitney, Proteolytically activated antibacterial toxins inhibit the growth of diverse Gram-positive bacteria. [Preprint] (2025). Available at: http://biorxiv.org/lookup/doi/10.1101/2025.04.13.648598 [Accessed 23 July 2025].

40. O. Goldbeck, D. Weixler, B. J. Eikmanns, C. U. Riedel, In Silico Prediction and Analysis of Unusual Lantibiotic Resistance Operons in the Genus Corynebacterium. Microorganisms 9, 646 (2021).

41. T. Argov, L. Rabinovich, N. Sigal, A. A. Herskovits, An Effective Counterselection System for Listeria monocytogenes and Its Use To Characterize the Monocin Genomic Region of Strain 10403S. Appl Environ Microbiol 83, e02927–16 (2017).

42. R. Simon, U. Priefer, A. Pühler, A Broad Host Range Mobilization System for In Vivo Genetic Engineering: Transposon Mutagenesis in Gram Negative Bacteria. Nat Biotechnol 1, 784–791 (1983).

43. L. M. Shull, et al., Analysis of genetic requirements and nutrient availability for *Staphylococcus aureus* growth in cystic fibrosis sputum. mBio 16 (2025).

44. C. F. Schuster, S. A. Howard, A. Gründling, Use of the counter selectable marker PheS* for genome engineering in Staphylococcus aureus. Microbiology (Reading) 165, 572–584 (2019).

45. I. R. Monk, T. P. Stinear, From cloning to mutant in 5 days: rapid allelic exchange in Staphylococcus aureus. Access Microbiol 3, 000193 (2021).

46. I. R. Monk, J. J. Tree, B. P. Howden, T. P. Stinear, T. J. Foster, Complete Bypass of Restriction Systems for Major Staphylococcus aureus Lineages. mBio 6, e00308–00315 (2015).

47. R. A. Forsyth, et al., A genome-wide strategy for the identification of essential genes in *Staphylococcus aureus*. Molecular Microbiology 43, 1387–1400 (2002).

48. A. R. Wattam, et al., Improvements to PATRIC, the all-bacterial Bioinformatics Database and Analysis Resource Center. Nucleic Acids Res 45, D535–D542 (2017).

49. R. D. Olson, et al., Introducing the Bacterial and Viral Bioinformatics Resource Center (BV-BRC): a resource combining PATRIC, IRD and ViPR. Nucleic Acids Research 51, D678– D689 (2023).

50. J. Abramson, et al., Accurate structure prediction of biomolecular interactions with AlphaFold 3. Nature 630, 493–500 (2024).

51. L. Holm, A. Laiho, P. Törönen, M. Salgado, DALI shines a light on remote homologs: One hundred discoveries. Protein Sci 32, e4519 (2023).

52. L. P. Kozlowski, IPC – Isoelectric Point Calculator. Biology Direct 11, 55 (2016).

